# Integrative Single-Cell RNA and ATAC Sequencing Reveals the Impact of Chronic Cigarette Smoking on Lung Epithelial Responses to Influenza and Hyperoxia

**DOI:** 10.1101/2025.04.06.647429

**Authors:** Pei-Chun Cha, Zhenyang Zou, Jessica Nouws, Charles S Dela Cruz, Lokesh Sharma, Xiting Yan, Maor Sauler

## Abstract

Cigarette smoke (CS) is a significant risk factor for developing acute respiratory distress syndrome (ARDS), but the cellular and molecular mechanisms linking smoking to ARDS susceptibility remain unclear. Our goal was to improve our understanding of these mechanisms. To address this, we established a mouse model comparing long-term CS exposure to non-smoking controls, examining responses to influenza infection and hyperoxia-induced lung injury. The mice were divided into six groups (n=1 per group): control (CON), cigarette smoke (CS), hyperoxia (HYP), influenza infection (FLU), cigarette smoke plus hyperoxia (CS+HYP), and cigarette smoke plus influenza infection (CS+FLU). Single-cell RNA sequencing (scRNA-seq) and single-cell ATAC sequencing (scATAC-seq) were performed on lung tissues. Quality control analysis using Seurat (v5.1.0) and Signac (v1.13.1) retained 78,402 cells from scRNA-seq and 84,144 cells from scATAC-seq, with 32,305 matched cells identified across both datasets. Differential gene expression analysis revealed significant smoking-associated alterations in cellular responses to influenza infection and hyperoxia exposure. Pathway enrichment indicated heightened immune responses, inflammatory signaling, and cellular survival pathways in smoking-exposed animals. Integration of scRNA-seq and scATAC-seq identified key transcription factors (TFs), including those involved in immune regulation, tissue repair, and chromatin remodeling mediating these responses. Overall, this study underscores the role of chronic cigarette smoke exposure in exacerbating pathways critical to ARDS pathogenesis, providing potential targets for therapeutic intervention.

## Introduction

Acute respiratory distress syndrome (ARDS) is a life-threatening condition characterized by diffuse alveolar damage, increased vascular permeability, and severe hypoxemia. It arises from a range of direct and indirect insults, including pneumonia, sepsis, aspiration, and trauma, with mechanical ventilation often exacerbating lung injury. While ARDS can develop in response to various triggers, underlying risk factors such as advanced age, chronic lung disease, and genetic predisposition influence disease susceptibility and severity. Despite advances in critical care, mortality remains high, and effective targeted therapies are lacking, underscoring the need for a deeper understanding of its pathophysiology [1].

Cigarette smoke (CS) exposure has been consistently associated with an increased risk of ARDS across multiple patient populations, including those with trauma, sepsis, and transfusion-related lung injury. Both active and passive smoking have been implicated, suggesting that even low levels of exposure can contribute to susceptibility. Epidemiological studies indicate that smokers who develop ARDS tend to be younger and may exhibit lower illness severity compared to nonsmokers, despite their increased risk of lung injury [2]. This paradox suggests that smoking may induce a state of baseline vulnerability, lowering the threshold for ARDS development upon subsequent insults. Moreover, CS exposure has been linked to an exaggerated inflammatory response in experimental models of lung injury, further supporting its role in ARDS pathogenesis [3]. However, while the association between smoking and ARDS risk is well-documented, the precise biological mechanisms remain incompletely understood, necessitating further investigation.

Chronic CS exposure disrupts lung homeostasis by inducing oxidative stress and alveolar-capillary barrier damage, which increases susceptibility to lung injury and pulmonary edema [4]. These changes, along with dysregulated inflammation and impaired fluid clearance, can exacerbate lung damage and heightens the risk of ARDS upon secondary insult. Beyond its direct cytotoxic effects, CS has been shown to induce epigenetic modifications that may contribute to ARDS susceptibility. Exposure to CS alters chromatin accessibility, DNA methylation, and histone modifications, leading to long-lasting changes in gene expression [5–7]. These epigenetic alterations can impair critical pathways involved in immune regulation, epithelial barrier function, and lung repair, ultimately predisposing individuals to a heightened inflammatory response upon injury.

Advances in single-cell sequencing provide an opportunity for evaluating epigenetic changes at single-cell resolution. In addition to gene expression, chromatin accessibility plays a key role in cellular responses to injury and disease by regulating transcriptional programs that contribute to lung pathology. Single-cell ATAC sequencing (scATAC-seq) enables genome-wide mapping of open chromatin regions, allowing for the study of epigenomic regulation in disease states. In this study, we applied single-cell RNA sequencing (scRNA-seq) and single-cell ATAC sequencing (scATAC-seq) to investigate how CS influences gene expression and chromatin accessibility in acute lung injury models. Using hyperoxia and acute viral infection models, we examined distinct mechanisms of lung injury, including oxidative stress-induced epithelial damage and immune-driven inflammation. By integrating transcriptomic and epigenomic data, we identified regulatory pathways affected by smoking, influenza, and hyperoxia. These findings provide a basis for future research into the molecular mechanisms underlying ARDS susceptibility and progression.

## Methods

### Animals

All animal procedures were approved by the Yale University Institutional Animal Care and Use Committee (IACUC) and conducted in accordance with the National Institutes of Health guidelines for the care and use of laboratory animals. AKR/J [mice were housed under standard laboratory conditions with free access to food and water. The mice were divided into six groups (n=1 per group): control (CON), cigarette smoke (CS), hyperoxia (HYP), influenza infection (FLU), cigarette smoke plus hyperoxia (CS+HYP), and cigarette smoke plus influenza infection (CS+FLU). Mice in the CS groups were exposed to CS for four months to mimic chronic smoking. The FLU groups were intranasally infected with 10 PFUs of H1N1 (PR8) strain of influenza in 50ul of PBS as described before (PMID: 39884393, PMID: 30110182) and the HYP groups were exposed to 100% oxygen for 48 hours to induce hyperoxia. At the end of the exposure periods, mice were euthanized and lung tissues were collected as previously described. Frozen lungs were snap frozen and stored in liquid nitrogen for further processing [8].

### Preparation of samples for sequencing libraries

Nuclei were isolated from human snap frozen lung tissue using the 10x genomics nuclei isolation kit, per manual (document number CG000505, revision A, May 2022) [9]. An aliquot of nuclei was stained with 0.4% trypan blue stain and DAPI and subsequently counted with the Thermo Fisher countess II FL. Samples with > 90% viability were used for downstream analysis. Approximately 10,000 nuclei per sample were loaded onto chip J and processed with the Chromium X Series to create to create Gel Beads-in-emulsion (GEMs). The next steps were carried out according to the manufacturer’s user guide (CG000338, revision G, September 2024) [10], including post-GEM-RT Dynabead cleanup and Pre-Amplification. Each sample was then split into two portions: one for ATAC library construction and the other for cDNA amplification and gene expression library construction. DNA from the ATAC libraries (scATAC libraries) and cDNA from the gene expression libraries (scRNA libraries) were assessed at two time points using the Agilent Bioanalyzer High Sensitivity Chip to ensure quality. Six scATAC libraries and six scRNA libraries were sequenced on the Novaseq 6000 platform, using 2 × 100 bp paired-end sequencing configuration.

### Preprocessing of single-cell RNA sequencing data

On average, 573,773,762 reads (s.d. =62,742,482) were sequenced for each scRNA library and 29,633 reads were sequenced per cell (s.d. = 11,739). The mean sequencing saturation was 78.43 ± 0.17%. The mean fraction of reads with a valid barcode was 91.32 ± 0.02% (Supplementary Table 1). The raw sequencing reads were preprocessed using the Cell Ranger pipeline (V7.1.0) with mouse genome (mm10) for cell calling and reads mapping to calculate the number of unique molecular identifier (nUMI) for each gene. Then the downstream analyses of the data were all performed using Seurat (v5.1.0) [11] unless otherwise described. Low-quality nuclei were removed (nFeatures > 200, RNA count > 200 and percentage of mitochondrial genes < 5%), and both debris and doublets were removed through iterative clustering. The filtered data was log-normalized and scaled. Dimensional reduction was conducted using principal component analysis (PCA) and Uniform Manifold Approximation and Projection (UMAP) was applied for data visualization [12,13]. Cell clustering was performed using the Louvain algorithm. For cell type annotation, a reference dataset [14] with cell type annotations was obtained and integrated with the collected data using the FindTransferAnchors function. The TransferData function was then applied to assign cell type annotation for each cell based on the cell type of the most similar cell from the reference data in the integrated data.

### Preprocessing of single-cell ATAC sequencing data

On average, 637,281,865 and 40,732 reads were sequenced for each scATAC library (s.d. = 118,797,857) and cell (s.d. = 6,760), respectively. The mean sequencing saturation for scATAC libraries was 66.91 ± 0.04% and the mean fraction of reads with a valid barcode was 97.51 ± 0.00% (Supplementary Table 2). The raw sequencing reads were preprocessed using the cellranger-atac pipeline (v2.1.0) with mouse genome annotation (mm10) from 10X Genomics to identify peaks and calculate the number of UMIs for each peak. The data was further processed using Seurat (v5.1.0) and its companion R package Signac (v1.13.1) [15]. Low-quality cells were removed from the aggregated scATAC-seq library (ATAC count > 400, nucleosome signal < 2 and TSS enrichment > 1). The filtered data was normalized using term-frequency inverse-document-frequency (TFIDF).

### Differentially expressed genes (DEGs) and differentially accessible peaks (DAPs) identification

To identify DEGs or DAPs between two given groups (CS vs. non-CS, influenza vs. non-influneza, hyperoxia vs. non-hyperoxia), we first filtered genes/peaks with > 0 nUMI in at least 7.5% of cells in all samples. Since there is only one sample per group, we let 𝑌_𝑖𝑗_ denote the nUMI for gene/peak 𝑗 in cell 𝑖. Then we assume that the nUMIs of gene/peak 𝑗 across all cells follows a negative binomial distribution: 𝑌_𝑖𝑗_ ∼ 𝑁𝐵(𝑆_𝑖_𝜇_𝑖𝑗_, 𝑑_𝑗_), where log(𝜇_𝑖𝑗_) = 𝛽_0𝑗_ + 𝛽_1𝑗_ ⋅ 𝑔𝑟𝑜𝑢𝑝_𝑖_, 𝑆_𝑖_ is the total number of UMIs in cell 𝑖, 𝑑_𝑗_ is the dispersion of the negative binomial distribution for gene/peak 𝑗. This model was fitted for each gene/peak separately to calculate the significance of 𝛽_1𝑗_. Genes with an FDR < 0.05 and an absolute log2 fold change (|β_1j|) > 0.5 were considered significant, while peaks were defined as significant based on a nominal p-value < 0.05 and an absolute log2 fold change (|β_1j|) > 0.5. To identify genes responding to the injury differently between smokers and non-smokers, i.e. the smoking-flu/hyperoxia interacted genes, we modified the model above by assuming log(𝜇_𝑖𝑗_) = 𝛽_0𝑗_ + 𝛽_1𝑗_ ⋅ 𝑠𝑚𝑜𝑘𝑒_𝑖_ + 𝛽_2𝑗_ ⋅ 𝑓𝑙𝑢_𝑖_ + 𝛽_3𝑗_ ⋅ 𝑠𝑚𝑜𝑘𝑒_𝑖_ ⋅ 𝑓𝑙𝑢_𝑖_. Significance of 𝛽_3𝑗_ was assessed and genes with an FDR < 0.05 and an absolute log2 fold change (|β_1j |) >0.5 were considered significant.

### Direct integration of scRNA-seq with scATAC-seq

To integrate the scRNA-seq and scATAC-seq data, we focused exclusively on cells that were jointly profiled in both datasets, identified through shared cell barcodes. The scATAC-seq peaks to genes using ChIPseeker (v1.38.0) [16]. DEGs identified from scRNA-seq were paired with DAPs identified from scATAC-seq that were assigned to the same gene. Pairs of DEG and DAP from the same gene were not considered as valid DEG-DAP pairs if the direction of regulation (either upregulation or downregulation) was inconsistent between the two datasets.

### Transcription factor (TF) integration of scRNA-seq with scATAC-seq and identification of top transcription factors (TFs) regulating the gene expression and chromatin accessibility

We incorporated the transcription factor binding sites (TFBS) information to detect potential TFs relevant to the dysregulation of DEGs. The promoter regions of DEGs were extracted using biomaRt (v 2.58.2) [17,18] and binding sites for all transcription factors within the promoter regions were identified within it using JASPAR2020 [19] database. We defined DEG-DAP pair by examining whether DAPs overlapped with TFBSs in DEG promoter regions. If a peak overlapped with a TFBS, the corresponding DEG and DAP were paired.

To prioritize key TFs of DEG-DAP pairs, we calculated a TF enrichment score as follows. The number of identified binding site of each TF was first divided by the total number of identified binding sites of all TFs in DEG to calculate the TF ratio before considering DAPs, denoted as 𝑇𝐹𝑅_𝑏𝑒𝑓𝑜𝑟𝑒_. Then we focused on the TFs in DEG-DAP pairs, and the TF ratio was recalculated if the number DEG-DAP pair was > 20, denoted as 𝑇𝐹𝑅_𝑎𝑓𝑡𝑒𝑟_. Finally, for each TF, we calculate a TF enrichment score using 𝑇𝐹𝑅_𝑎𝑓𝑡𝑒𝑟_/𝑇𝐹𝑅_𝑏𝑒𝑓𝑜𝑟𝑒_. The top 10 TFs with the largest TF enrichment score were selected, provided their absolute enrichment score was ≥ 0.5; if fewer than 10 TFs met this threshold, we selected only those that did. These enriched TFs are likely to play important regulatory roles, as their enhanced presence in DAP-associated regions indicates a stronger influence on the gene expression changes of the DEGs.

### Pathway enrichment analysis

The DEGs of CS, flu and hyperoxia, as well as the interacting genes identified from smoking-flu/hyperoxia interaction, were subjected to pathway enrichment analysis using Metacore. We used the enrichment analysis tool to identify the enriched pathways involving these genes. Based on a significant p-value < 0.05, graphical depictions of pathways were then generated.

## Results

### Cell populations identified in the scATAC-seq dataset using scRNA-seq as a reference

To assess the consequences of CS exposure on susceptibility to acute lung injury, we performed scRNA-seq and scATAC-seq on lung tissue from mice exposed to either CS or room air, followed by either hyperoxia or acute influenza infection. This experimental design allowed for the assessment of gene expression and chromatin accessibility changes associated with CS exposure in the context of acute lung injury (Fig. 1A). Cellular clusters were annotated based on the reference scRNAseq dataset (Fig. 1B, Left). Focusing on epithelial cells, we identified distinct canonical subtypes, including alveolar type 1 (AT1), alveolar type 2 (AT2), ciliated and secretory cells that expressed corresponding marker genes (Fig. 1B, Right bottom). Cell labels for scATAC-seq scATAC-seq were assigned by aligning barcodes to scRNA-seq data. After pooling all samples, we applied UMAP to visualize both datasets. The UMAP projections showed consistent clustering by cell type or condition, highlighting parallel patterns in transcriptional profiles and chromatin accessibility. (Fig. 1C).

**Figure 1.**
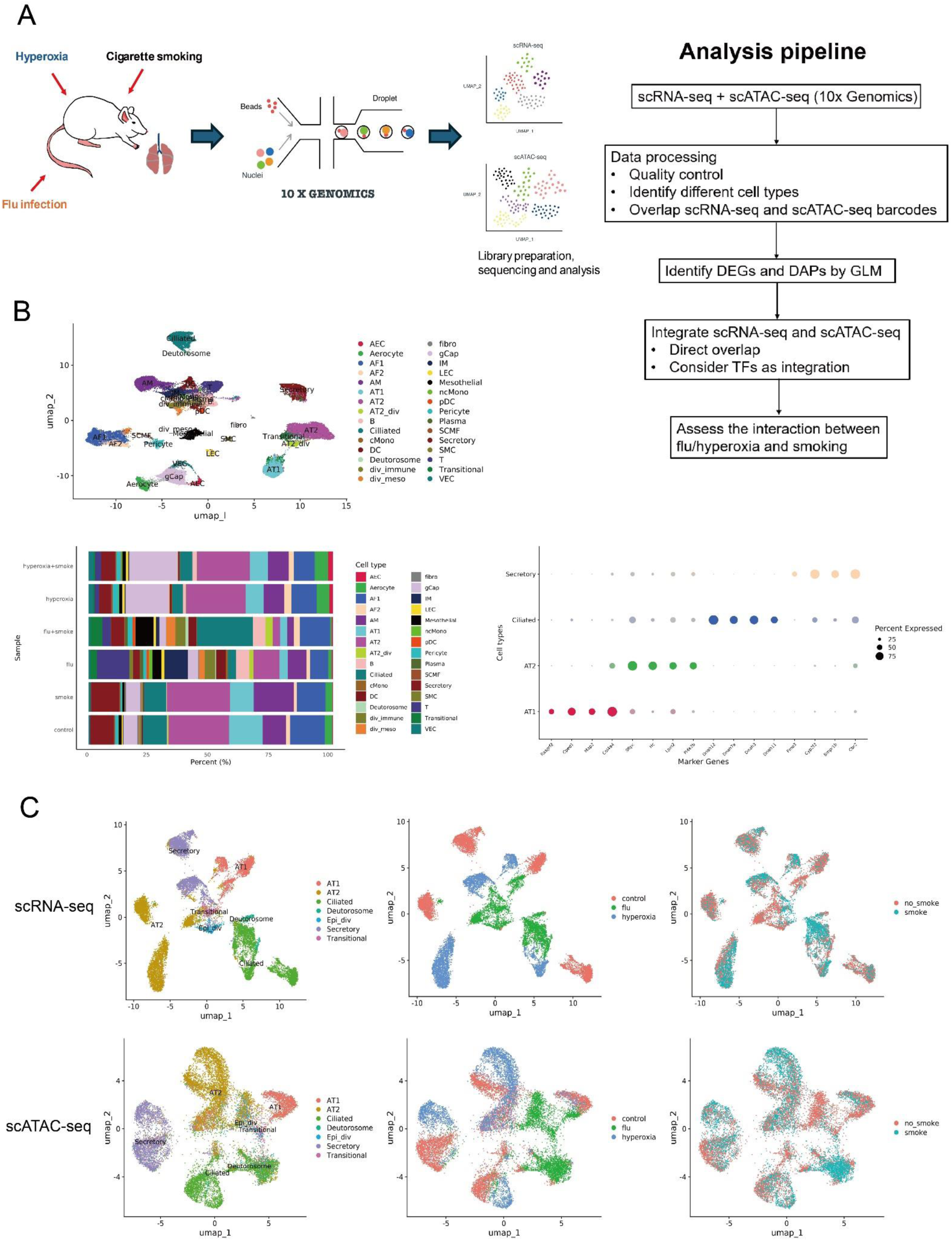
Single cell sequencing workflow and cell-type annotation of the single-cell ATAC sequencing (scATAC-seq) dataset using single-cell RNA sequencing (scRNA-seq) as a reference. (A) Graphical abstract of experimental methodology and analysis pipeline. Mice lung tissues were assessed with scRNA-seq and scATAC-seq (n=1/group). Abbreviations: DEGs, differentially expressed genes; DAPs, differentially accessible peaks; GLM, generalized linear model; TFs, transcription factors. (B) Left top: Uniform Manifold Approximation and Projection (UMAP) visualization of single cells classified by scRNA-seq data. Left bottom: Markers genes of epithelial cells. Right bottom: Proportions of cell types across each sample. (C) UMAP plots of scRNA-seq and scATAC-seq dataset, clustered by cell types, condition (control, influenza (flu), hyperoxia), and smoking status.

### Identification of DEGs and DAPs

We identified transcriptional and epigenetic alterations, characterized by DEGs and DAPs, within epithelial cell populations subjected to influenza infection and hyperoxic injury. (Supplementary Table 3). Pathway enrichment analysis of the DEGs highlighted distinct biological responses to these exposures. In influenza-infected samples, immune response pathways were predominantly activated, underscoring host defense mechanisms. These included BCR pathways, IL-3 signaling and IL-4 signaling. Under hyperoxia condition, we found significant activation of survival and developmental pathways, such as chemotaxis mediated by lysophosphatidic acid (LPA) signaling via GPCRs and the regulation of STK (Src family tyrosine kinases) signaling (Fig. 2).

**Figure 2.**
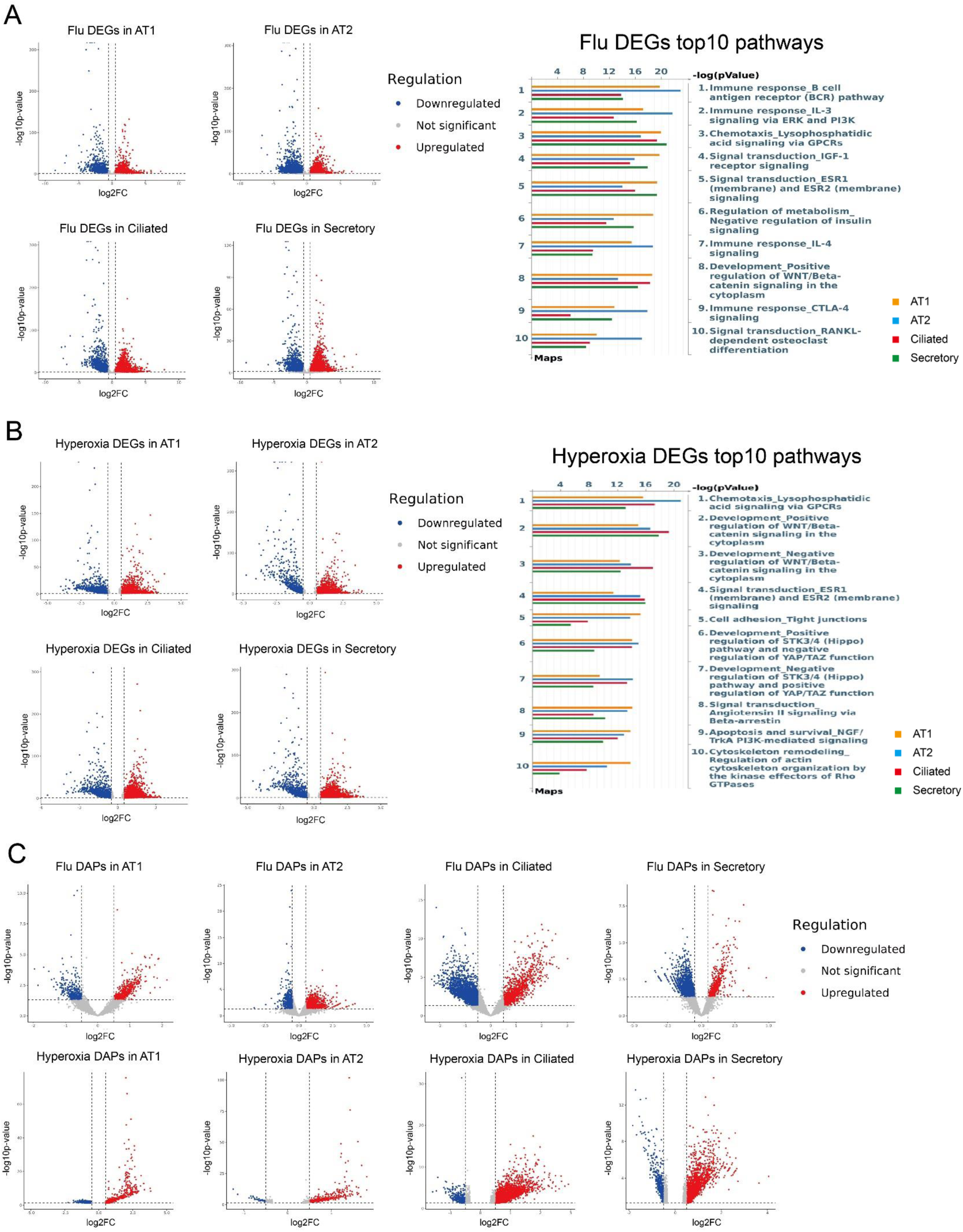
Differentially expressed genes (DEGs) from single-cell RNA sequencing (scRNA-seq) data and differentially accessible peaks (DAPs) from single-cell ATAC sequencing (scATAC-seq) under influenza infection and hyperoxia. (A) Left: Volcano plots displayed DEGs in response to influenza (flu) infection, showing upregulated and downregulated genes. Genes with significant upregulation were highlighted with red, and those with significant downregulation were highlighted with blue. Right: Top 10 enriched pathways from analysis of influenza (flu) upregulated DEGs. Pathway enrichment analysis highlighted biological processes significantly associated with influenza (flu)-affected genes. (B) Left: Volcano plots displayed the DEGs in response to hyperoxia, showing upregulated and downregulated genes. Genes with significant upregulation were highlighted with red, and those with significant downregulation were highlighted with blue. Right: Top 10 enriched pathways from analysis of hyperoxia upregulated DEGs. Pathway enrichment analysis highlighted biological processes significantly associated with hyperoxia-affected genes. (C) Top: Volcano plots displayed DAPs in response to influenza (flu) infection, showing upregulated and downregulated peaks. Peaks with significant upregulation were highlighted with red, and those with significant downregulation were highlighted with blue. Bottom: Volcano plots displayed DAPs in response to hyperoxia, showing upregulated and downregulated peaks. Peaks with significant upregulation were highlighted with red, and those with significant downregulation were highlighted with blue.

To investigate whether the changes in open chromatin regions were correlated with the changes in gene expression levels, we performed an integrative analysis of the scATACseq and scRNA-seq data sets in two ways. We performed direct integration, in which we identified DAP-DEG pairs based on genomic proximity and consistent regulation direction, while TF integration overlayed transcription factor binding site data to identify regulatory elements likely to mediate gene expression changes.

The numbers of DEG-DAP pairs were shown in Supplementary Table 3 and Fig. 3A. alveolar type 1 (AT1) cells, we identified 201 DEG-DAP pairs through direct integration under influenza conditions, while the number for hyperoxia were 162. For alveolar type 2 (AT2) cells, we found 533 DEG-DAP pairs associated with influenza, and the number under hyperoxia was 111. Ciliated cells exhibited 954 DEG-DAP pairs under influenza conditions and 637 under hyperoxia conditions. Additionally, in secretory cells, 329 DEG-DAP pairs were found, and the number associated with hyperoxia was 505. Further integration of TF information reduced the number of DEG-DAP pairs by slightly less than 50% (Fig. 3A, Supplementary Table 3). The heatmaps of DEGs identified through TF integration were shown in Fig. 3B. Although this approach identified fewer DEG-DAP pairs than direct integration, it provided deeper insights into regulatory mechanisms.

**Figure 3.**
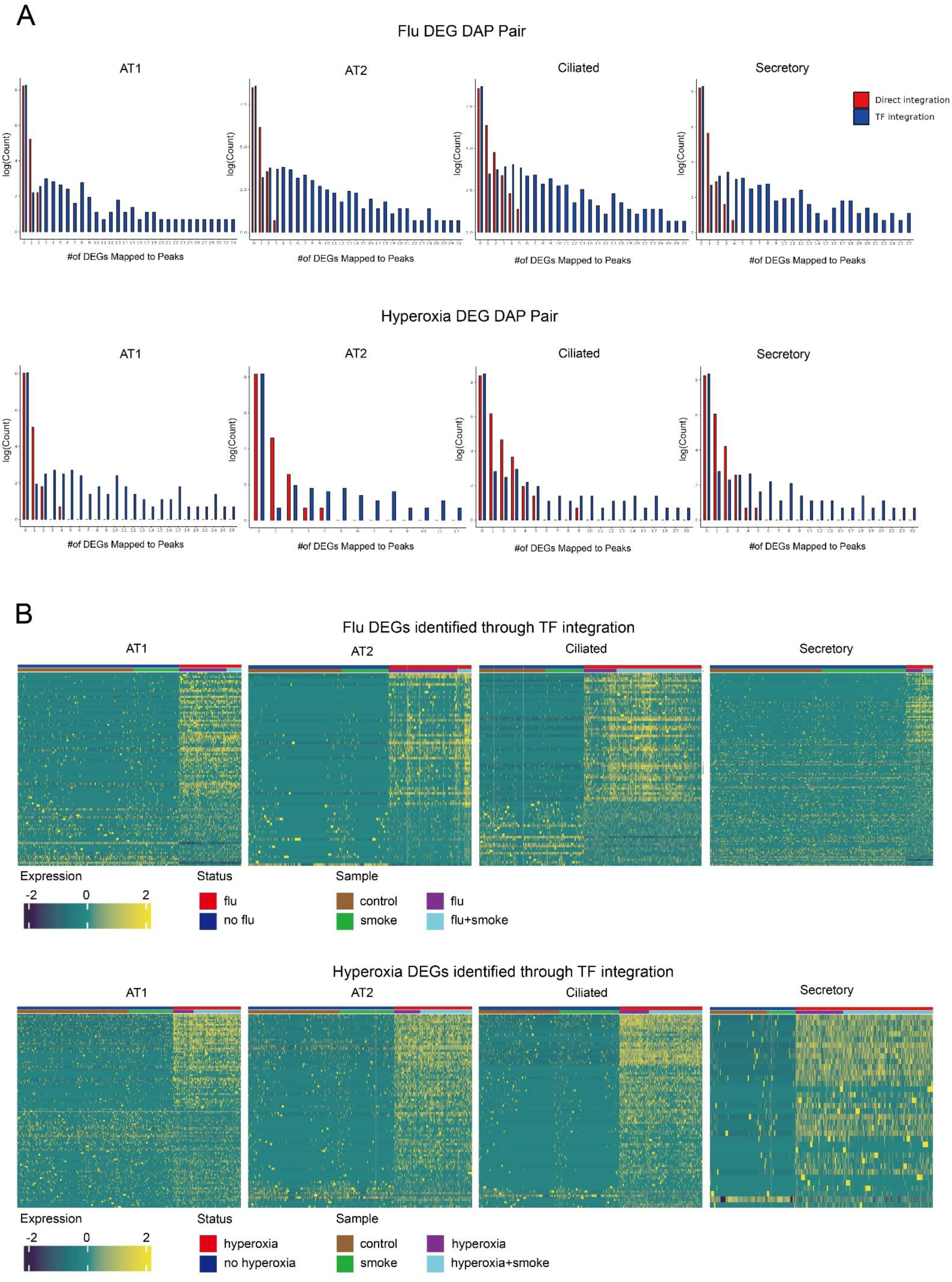
Integration of single-cell RNA sequencing (scRNA-seq) and single-cell ATAC sequencing (scATAC-seq). (A) Top: Histograms showing distribution of the number influenza (flu) differentially expressed genes (DEGs) linked per peak by direct and transcription factor (TF) integration. The X-axis represents the number of a specific DEG mapped to differentially accessible peaks (DAPs), while the Y-axis indicates the frequency of occurrences for each number. Bottom: Histograms showing distribution of the number hyperoxia DEGs linked per peak by direct and TF integration. The X-axis represents the number of a specific DEG mapped to DAPs, while the Y-axis indicates the frequency of occurrences for each number. (B) Top: Heatmaps displayed DEGs in response to influenza (flu) infection across four epithelial cell types (AT1, AT2, Ciliated, and Secretory). Bottom: Heatmaps showed hyperoxia-responsive DEGs in the same four epithelial cell types. Both analyses utilized TF integration to identify cell type-specific gene expression patterns associated with influenza (flu) infection and hyperoxia exposure.

### Potential transcription factors associated with DEGs of flu and hyperoxia

To further understand the regulatory role of transcription factors, we identified the specific TFs enriched amongst DEGs identified via the TF integration method (Fig. 4). For influenza-upregulated DEGs, immune-related TFs such as STAT family (e.g., STAT1, STAT3) and RFX7 were consistently identified across epithelial cell types. Interestingly, members of the KLF (e.g., KLF2, KLF3, KLF5, KLF10, KLF11, KLF16) family were prominent among influenza-downregulated DEGs. Among the TFs of hyperoxia-upregulated DEGs, AT2 and secretory cells exhibited a unique enrichment of JUN and FOS, while the basic helix-loop-helix (bHLH) family (e.g., BHLHA15 and MSC) was increased in AT1 and ciliated cells.

**Figure 4.**
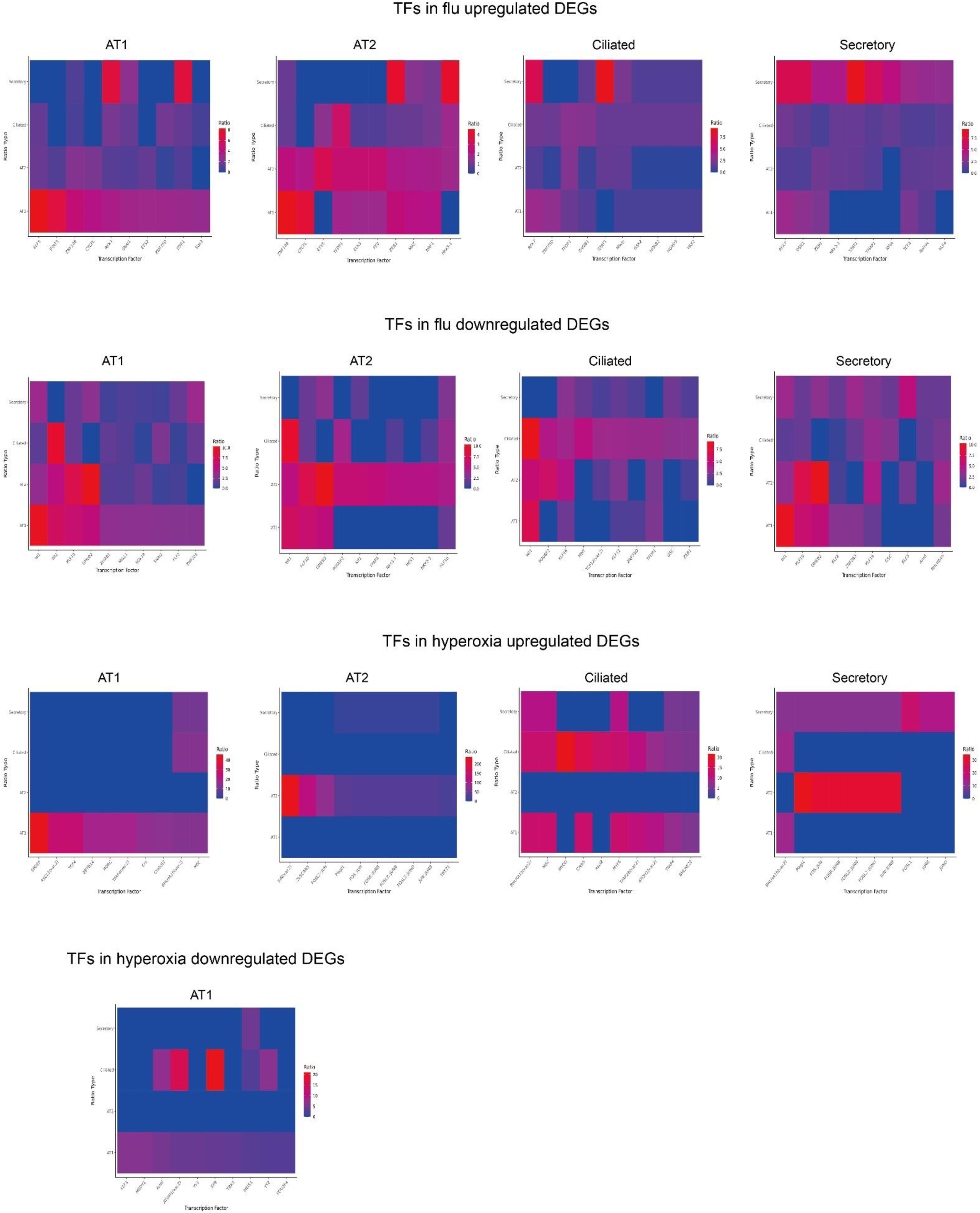
The top 10 transcription factors (TFs) in differentially expressed gene-differentially accessible peak (DEG-DAP) pairs for each cell type under influenza (flu) infection and hyperoxia conditions after TF integarion of single-cell RNA sequencing (scRNA-seq) and single-cell ATAC sequencing (scATAC-seq). Top 10 transcription factors (TFs) for each cell type, selected based on the highest TF abundance ratio before and after integration with DAP data, with a minimum ratio threshold of 0.5 for consideration. Top: Heatmap representing transcription factor expression for DEG-DAP pairs in response to infleunza (flu) exposure, categorized by flu-upregulated DEGs (increased gene expression and chromatin accessibility under influenza (flu) infection) and flu-downregulated DEGs (decreased gene expression and chromatin accessibility under influenza (flu) infection). Rows represent four epithelial cell types, and columns correspond to the top 10 transcription factors. Bottom: Heatmap showing transcription factor expression for DEG-DAP pairs under hyperoxia conditions, categorized by hyperoxia-upregulated DEGs (increased gene expression and chromatin accessibility under hyperoxia exposure) and hyperoxia-downregulated DEGs (decreased gene expression and chromatin accessibility under hyperoxia exposure). Rows represent four epithelial cell types, and columns correspond to the top 10 transcription factors for each cell type.

### The responses to influenza infection and hyperoxia affected by CS

The impact of smoking on gene express was far reduced when compared to the impact of influenza and hyperoxia on gene expression, however, we did identify discreet changes (Fig. 5, Supplementary Table 4). Pathway analysis of CS-associated DEGs revealed significant changes associated with proteostasis, heat shock proteins, neurophysiologic processes, WNT signaling, epithelial-to-mesenchymal transition, and chemotaxis.

**Figure 5.**
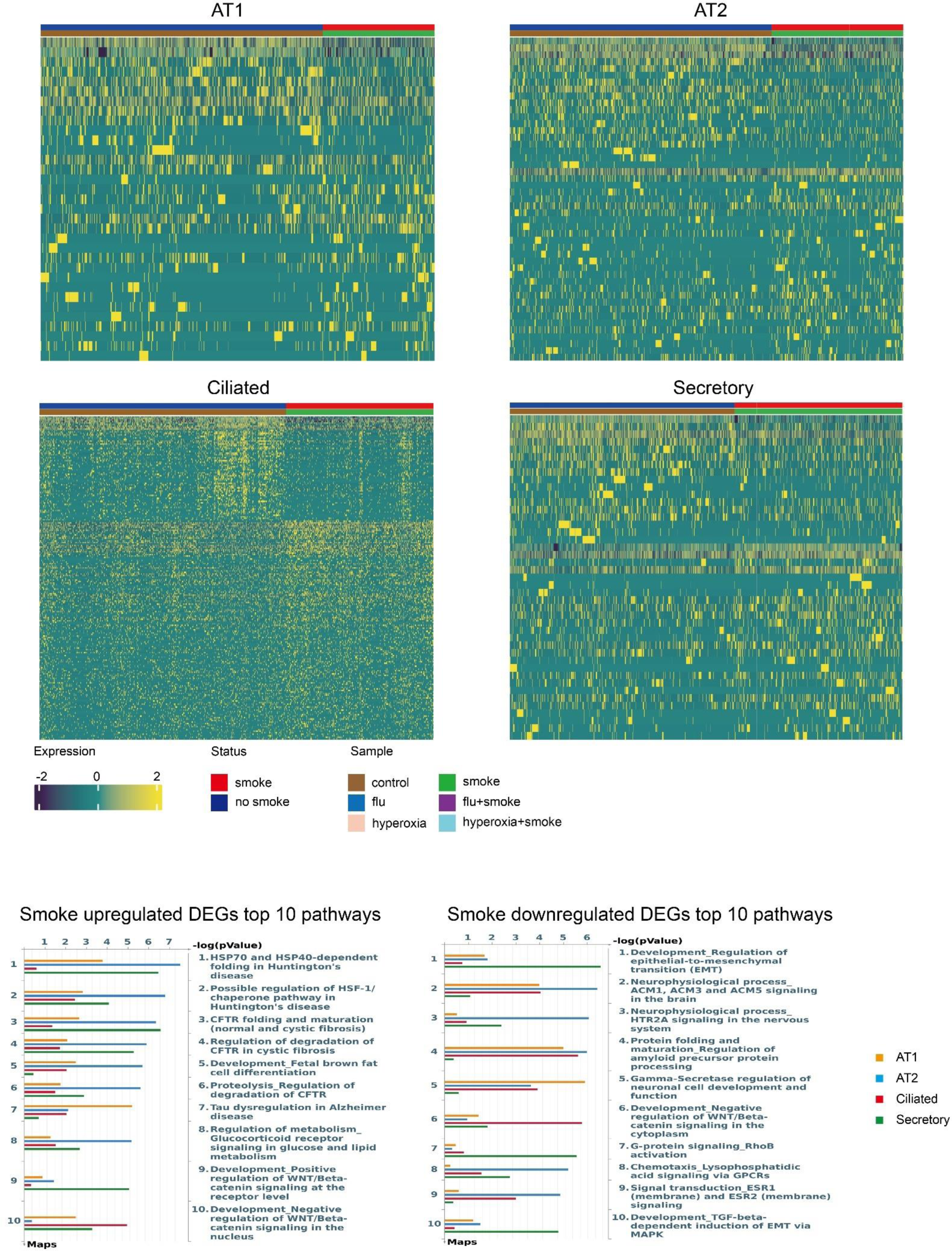
Differentially expressed genes (DEGs) from single-cell RNA sequencing (scRNA-seq) data in response to smoking. Top: Heatmap showing DEGs across four epithelial cell types (AT1, AT2, Ciliated, and Secretory) in response to smoking. Bottom: Pathway enrichment analysis of smoking-associated DEGs. Pathways enriched for positively regulated DEGs are displayed on the left, while pathways enriched for negatively regulated DEGs are displayed on the right.

We then sought to determine the impact of smoking on the cellular response to influenza infection. First, we examined gene expression changes only. We found increased expression of genes related to IL-3 mediated immune responses, ERK and PI3K signaling, mast cell activation, and VEGF signaling in that in mice exposed to CS and influenza vs. mice exposed to influenza alone. Conversely, smoking further suppressed genes already decreased following influenza infection, especially those involved in neurophysiological signaling, gamma-secretase regulation, and tight junction integrity. Smoking also impaired pathways that were increased in influenza-exposed mice, such as Hippo-YAP/TAZ, mTOR signaling, and FAK1, many of which are critical for epithelial homeostasis, repair, and barrier function. These changes suggest that smoking exacerbates the inflammatory and injurious transcriptional programs triggered by influenza while dampening key reparative and developmental responses, particularly in epithelial subsets like AT2 and Ciliated cells that are essential for lung regeneration and mucociliary clearance.

We also evaluated the impact of smoking on genes induced or suppressed by hyperoxia (Fig. 7, Supplementary Table 6). Smoking enhances gene expression in hyperoxia for pathways related to apoptosis and cell survival (e.g., NGF/TrkA and PI3K signaling), protein folding, cytoskeletal remodeling, and neural signaling, especially notable in AT2 and Ciliated cells. Similarly, CS exposure was associated with a further reduction in the expression of genes involved in metabolism, synaptic signlaing, insulin and IGF signaling, and cytokine pathways (e.g., IL-2, IL-4, and IL-6). Smoking also blunted genes that were increased following hyperoxia exposure, reated to tight junction integrity, WNT/β-catenin signaling, and immune regulation (e.g., IL-6/IL-8 cascades). This suppression suggests impaired epithelial barrier maintenance and immune crosstalk, likely compromising repair and regeneration, particularly in AT1 and Ciliated cells. Finally, smoking reduces the repression of certain hyperoxia-downregulated genes tied to metabolism (e.g., mTORC2, glucocorticoid signaling), development (Hippo/YAP-TAZ, ERBB), and Notch signaling. Collectively, these findings suggest that smoking prevents the normal resolution or adaptation to oxidative stress in epithelial cells.

**Figure 6.**
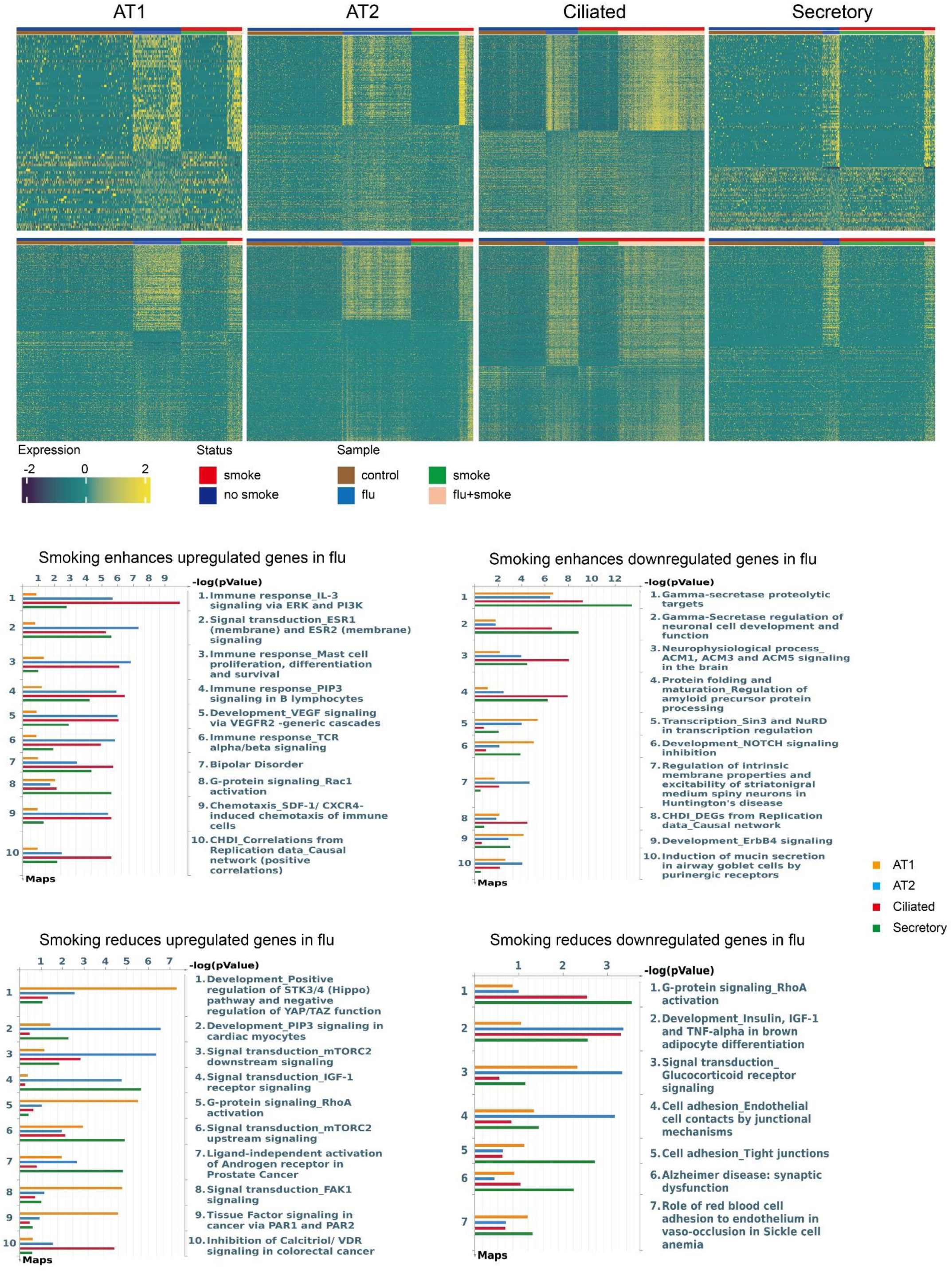
Single-cell RNA sequencing (scRNA-seq) analysis of infleunza (flu) infection in response to smoking. Top: Heatmaps showing positively interacting influenza (flu)-smoking genes in the upper panels, where smoking enhances the expression of influenza (flu)-upregulated genes or suppresses flu-downregulated genes, and negatively interacting flu-smoking genes in the lower panels, where smoking suppresses influenza (flu)-upregulated genes or enhances flu-downregulated genes, across four epithelial cell types (AT1, AT2, Ciliated, and Secretory). Bottom: Pathway enrichment analysis for positively and negatively smoking-infleunza (flu) interacting genes.

**Figure 7.**
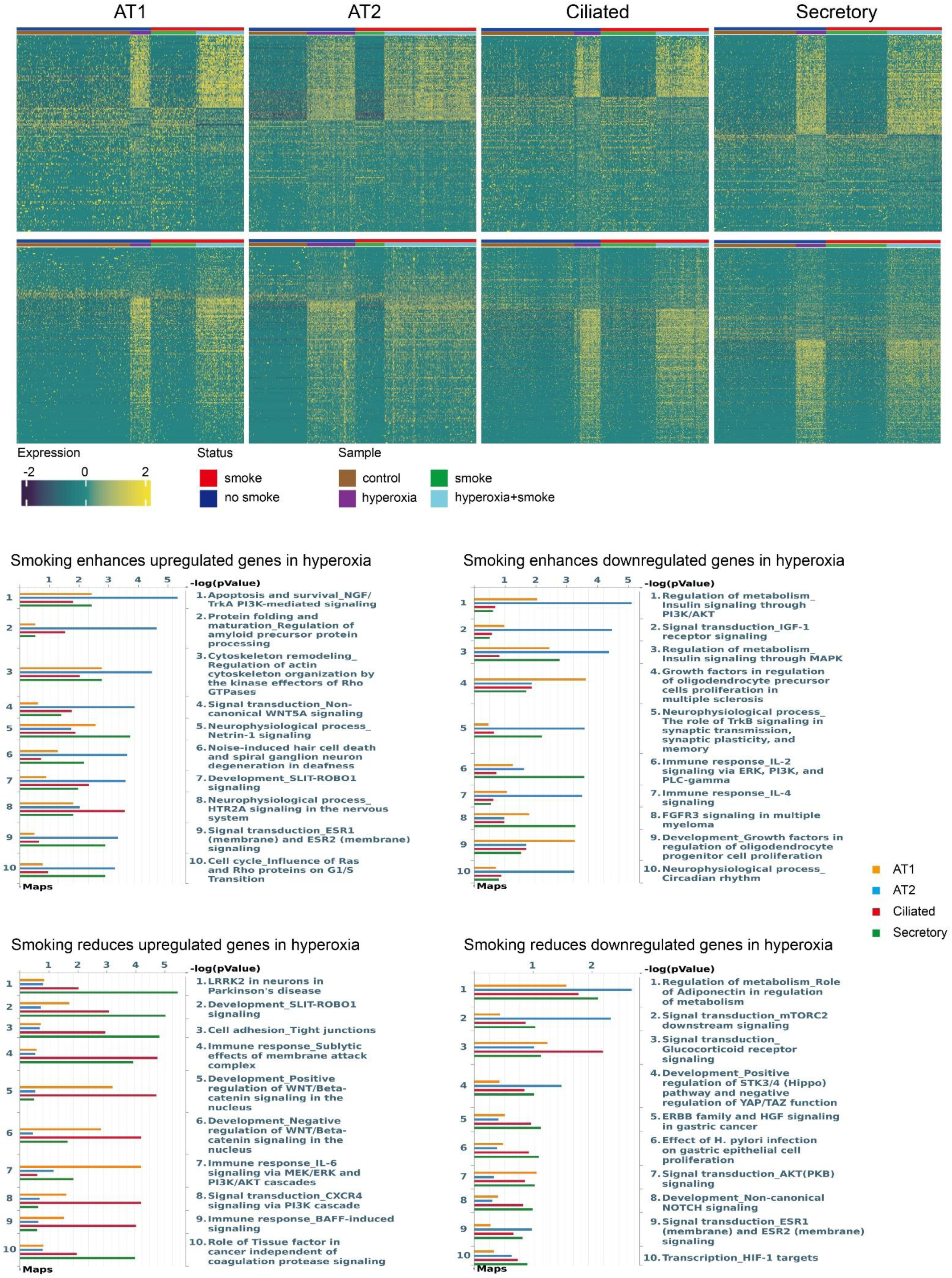
Single-cell RNA sequencing (scRNA-seq) analysis of hyperoxia in response to smoking. Top: Heatmaps showing positively interacting hyperoxia-smoking genes in the upper panels, where smoking enhances the expression of hyperoxia-upregulated genes or suppresses hyperoxia-downregulated genes, and negatively interacting hyperoxia-smoking genes in the lower panels, where smoking suppresses hyperoxia-upregulated genes or enhances hyperoxia-downregulated genes, across four epithelial cell types (AT1, AT2, Ciliated, and Secretory). Bottom: Pathway enrichment analysis for positively and negatively smoking-hyperoxia interacting genes.

Next, we determined the interaction between smoking and influenza on DEG-DEP pairs determined through direct integration (Supplemental Fig. 1). Because the number of gene-peak pairs in the smoking-hyperoxia interaction, and AT1 and secretory smoking-influenza interaction was fewer than 20, we excluded them from detailed analysis. Our findings suggest both gene-level (Fig. 6) and integrated multiomic analyses (Supplemental Fig. 1) Specifically, CS exposure further increased the expression of genes associated with immune and inflammatory responses, including IL-3 signaling, CXCR4 chemotaxis, and ERK/PI3K cascades, particularly in AT2 and Ciliated epithelial cells. CS exposure further reduced the expression of genes involved in neurophysiological and structural maintenance pathways, such as amyloid processing, gamma-secretase regulation, and NuRD-mediated transcriptional regulation, with pronounced effects in Ciliated cells. In addition, CS exposure blunted the increased expression of genes linked to epithelial repair and regeneration, including WNT/β-catenin signaling, NOTCH signaling, and adhesion, while also attenuating the normal downregulation of developmental and differentiation-related genes, collectively suggesting that CS mpairs epithelial homeostasis and injury recovery.

In AT2 cells, TF integration revealed that CS exposure enhanced the enrichment of several ETS family members—including ETV, ERG, ELK1, and ELK3—along with components of MAPK signaling such as ERK, suggesting reinforcement of pro-inflammatory and stress-responsive transcriptional programs. Conversely, among influenza-upregulated TFs, CS exposure reduced the activity of key epithelial regulators, including GATA5, SOX family members, STAT2, NKX6-3, and members of the POU family, all of which are associated with epithelial differentiation and barrier maintenance. Within influenza-downregulated genes, CS further suppressed the activity of SOX, ETV, and ELF transcription factors in AT2 cells, and led to pronounced reductions in HOX family TFs in both AT2 and Ciliated cells. These findings suggest that CS skews epithelial transcriptional regulation by enhancing stress-related TF activity while broadly suppressing developmental and differentiation-associated TF networks. (Fig. 8, Supplementary Table 7).

**Fig. 8.**
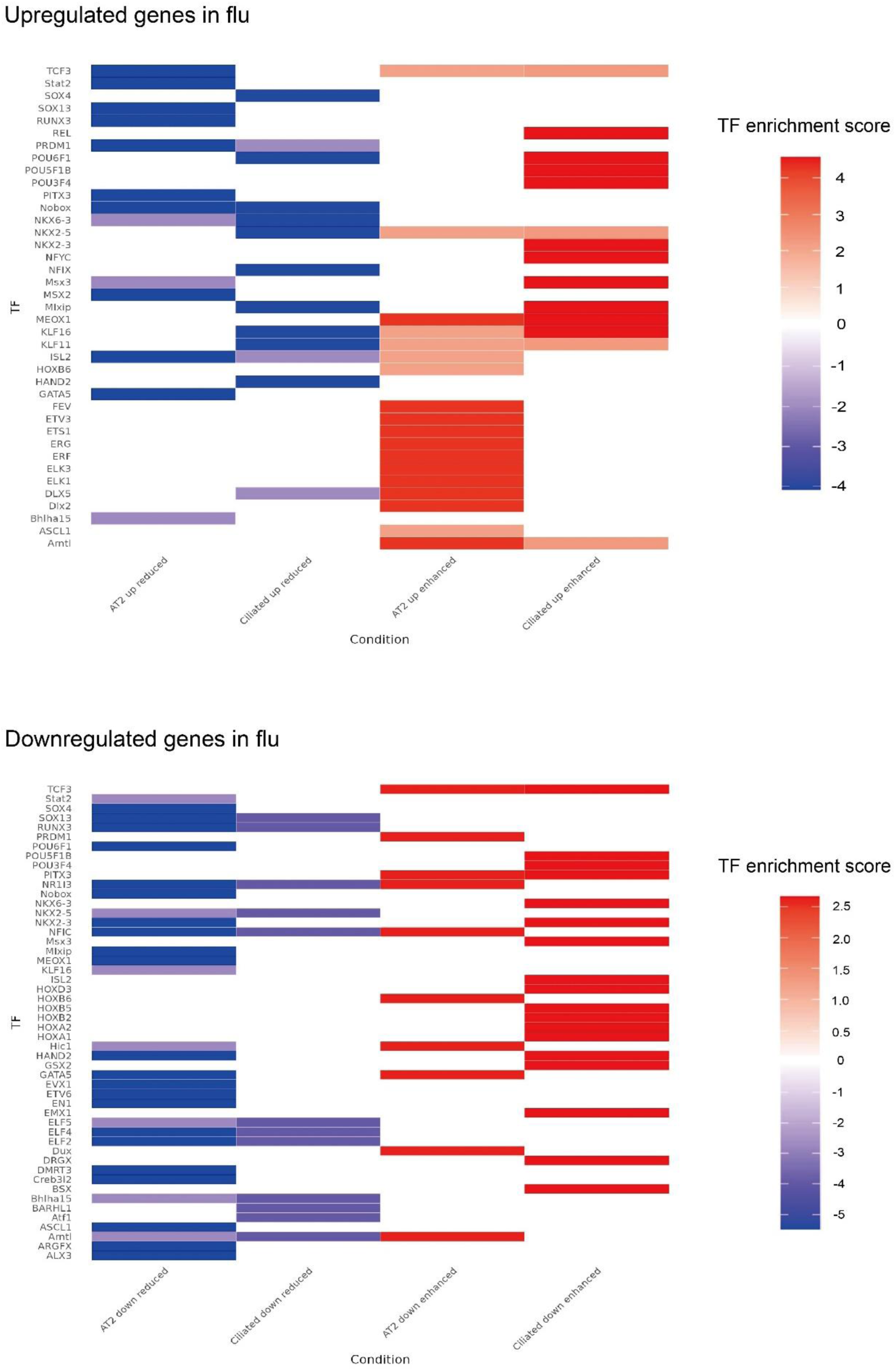
The top transcription factors (TFs) in differentially expressed gene-differentially accessible peak (DEG-DAP) pairs for each cell type under smoking-infleunza (flu) interaction condition after TF integarion of single-cell RNA sequencing (scRNA-seq) and single-cell ATAC sequencing (scATAC-seq). Top TFs for each cell type, selected based on the highest TF abundance ratio before and after integration with DAP data, with a minimum ratio threshold of 0.5 for consideration. Rows represent epithelial cell types (AT2, Ciliated) and different smoking-influenza (flu) interaction conditions: up reduced (smoking suppresses flu-upregulated genes), up enhanced (smoking enhances flu-upregulated genes), down reduced (smoking enhances flu-downregulated genes), and down enhanced (smoking suppresses flu-downregulated genes). Columns represent the top 10 TFs, combined across different conditions and cell types in a single plot.

## Discussion

In this study, we characterized gene expression and chromatin accessibility in epithelial cells to explore how CS exposure influences the subsequent response to influenza or hyperoxia. Through integrative single-cell RNA and ATAC sequencing, we uncover a transcriptional landscape in which CS exposure amplifies inflammatory responses while blunting epithelial repair and differentiation programs in response to influenza infection and hyperoxia. This dual effect is mediated, at least in part, by altered activity of key TF families that orchestrate epithelial homeostasis and regeneration in the lung.

Previous studies in mice have demonstrated that CS exposure increases susceptibility to ALI. Sakhatskyy et al. developed a “double-hit” mouse model to investigate how CS exposure primes the lungs for heightened ALI upon subsequent lipopolysaccharide (LPS) challenge. Both acute (3-hour) and subacute (3-week) CS exposures enhanced LPS-induced lung injury in C57BL/6 and AKR mouse strains, with the AKR mice, which are more susceptible to CS, exhibited greater susceptibility to LPS exposure following CS exposure, with a more pronounced change in gene expression changes [20–22]. However, the mechanisms underlying this increased susceptibility remain uncertain.

Using our multiomic approach, we identified epigenetic consequences of smoking that may impact suscpetibiility to lung injury. Among the most striking findings is the suppression of SOX family transcription factors, notably SOX4 and SOX13, in the context of smoking-influenza interactions. These TFs are critical regulators of epithelial identity and epithelial-mesenchymal plasticity [23]. Their reduced activity, alongside diminished expression of canonical WNT/β-catenin and NOTCH signaling components, suggests a disruption of cell fate commitment and regenerative capacity in alveolar and airway epithelial cells [24]. WNT/β-catenin and NOTCH signaling have well-documented roles in AT2 cell proliferation, differentiation into AT1 cells, and in maintaining barrier integrity during and after injury. The blunting of these pathways likely contributes to the impaired epithelial repair and persistent barrier dysfunction observed in CS-exposed lungs following injury.

In parallel, we observed a marked reduction in the activity of HOX transcription factors—including HOXA1, HOXB5, and HOXD3, which are essential for lung morphogenesis and regional epithelial patterning. Prior studies have implicated HOX gene dysregulation in the pathogenesis of chronic obstructive pulmonary disease (COPD) and emphysema, where disrupted spatial identity and epithelial remodeling are prominent [25]. Our data extend these findings to the acute injury setting, suggesting that HOX suppression underlies a failure of proper epithelial regeneration and structural reconstitution after insult.

Conversely, CS exposure enhances the activity of pro-inflammatory ETS family TFs, including ETS1, ERG, ELK1, and ELK3. These factors are known mediators of cytokine production, chemokine signaling, and oxidative stress responses, and their increased activity aligns with the upregulation of ERK/PI3K cascades, CXCR4-mediated chemotaxis, and VEGF signaling observed in CS-exposed, influenza-infected epithelial cells [26–28]. This exaggerated inflammatory transcriptional program may drive sustained immune cell recruitment, tissue injury, and ineffective resolution, consistent with the enhanced susceptibility of smokers to ALI and the chronic inflammation observed in smoking-related lung disease.

Together, these findings suggest that CS reprograms the epithelial transcriptional network to favor injury amplification over repair. The coordinated suppression of SOX, WNT, NOTCH, and HOX-driven pathways, alongside the activation of ETS-mediated inflammatory programs, represents a shift toward a maladaptive epithelial state, one characterized by impaired regeneration, sustained inflammation, and increased vulnerability to secondary insults. These insights provide a mechanistic foundation for the heightened risk of ARDS in smokers and point toward transcriptional regulators as potential targets for therapeutic intervention in smoke-exacerbated lung injury.

We also found an interesting impact of smoking on hyperoxia related gene expression. For hyperoxia-upregulated DEGs, both AT2 and secretory cells displayed an enrichment of JUN and FOS. These TFs form the AP-1 complex, which regulates genes involved in cellular responses to stress and influences susceptibility to hyperoxia-induced cell death in murine lung epithelial cells [29–31].

While our study provides important mechanistic insights into how CS reprograms the lung epithelial response to injury, several limitations should be acknowledged. First, the sample size was necessarily constrained by the complexity and cost of single-cell multiomic profiling. With only one mouse per exposure group, the data do not capture inter-animal variability, and thus the results should be interpreted as proof-of-principle rather than definitive population-level effects. Additionally, our study focused on a single post-injury time point. This limits our ability to track dynamic transcriptional or epigenetic changes over time and precludes resolution of early injury versus late reparative responses. More broadly, multiomic integration remains a developing field with significant technical and analytical challenges. Although we leveraged direct and transcription factor-based integration to link chromatin accessibility to gene expression, these methods rely on assumptions about regulatory proximity and TF binding site annotation that may not fully capture the complexity of chromatin architecture or cell-type-specific regulation. Moreover, the sparsity and noise inherent to scATAC-seq data can limit sensitivity, particularly in rare cell populations. However, a strength of the study was that we were broadly able to replicate stereotypical responses to injury. For example, after influenza infection, we identified adaptive immune signaling pathways, and TRF activity related to STAT family and RFX7 activity across various epithelial cell types, while members of the KLF transcription factors (Krüppel-like factors) were prominently associated with influenza-downregulated DEGs [32]. In pulmonary epithleial cells exposed to hyperoxia, we found DEGs enriched for activation of LPA signaling and DEGs associated with both positive and negative STK and signaling pathways, pathways also dentified in other studies involving hyperoxia [33–35].

Despite the limitations of ours study, our findings highlight how CS reshapes the transcriptional and regulatory landscape of epithelial cells during acute lung injury. Larger cohorts, temporal resolution, and further refinement of integrative models will be essential to fully realize the potential of multiomic approaches in dissecting complex lung disease mechanisms.

## Contributors

M.S. conceived of and designed the study. J.N., C.D.C. and L.S. collected the samples and performed the experiments. P.C., Z.Z., X.Y. and M.S. analyzed the data, P.C., Z.Z., X.Y. and M.S. interpreted the results and wrote the manuscript.

## Declaration of interests

We declare that we have no conflicts of interests.

## Acknowledgement

The authors thank the Yale Center for Research Computing for the support in High Performance Computing.

## Funding

MS is supported by NIH grants R01HL155948, R21HL173512 and DOD grants W81XWH2210629,XY is supported by R01LM014087

## Notes

### Competing Interest Statement

The authors have declared no competing interest.

