## Supplementary files for "Integrative Single-Cell RNA and ATAC Sequencing Reveals the Impact of Chronic Cigarette Smoking on Lung Epithelial Responses to Influenza and Hyperoxia"

| **Sample** | **Total RNA reads** | **Mean RNA reads per cell** | **Sequencing saturation** | **Valid barcodes** |
| --- | --- | --- | --- | --- |
| **1** | 682,094,113 | 26,896 | 92.50% | 90.40% |
| **2** | 633,144,396 | 35,480 | 95.70% | 88.80% |
| **3** | 502,103,364 | 11,464 | 57.80% | 94.20% |
| **4** | 547,525,062 | 21,461 | 51.20% | 91.90% |
| **5** | 531,751,818 | 48,942 | 88.50% | 91.20% |
| **6** | 546,023,817 | 33,556 | 84.90% | 91.40% |
| **mean** | 573,773,762 | 29,633 | 78.43% | 91.32% |
| **stdev** | 62742481.69 | 11738.82542 | 0.173520092 | 0.01625235 |

**Supplementary Table1 Summary of single-cell RNA sequencing (scRNA-seq).** Summarizes the scRNA-seq data for six samples, showing an average of 573.8 million total RNA reads, 29,633 mean reads per cell, 78.43% sequencing saturation, and 91.32% valid barcode rates.

| **Sample** | **Total ATAC reads** | **Mean ATAC reads per cell** | **Sequencing saturation** | **Valid barcodes** |
| --- | --- | --- | --- | --- |
| **1** | 676844825 | 33833.7828 | 69.05% | 97.39% |
| **2** | 715285783 | 42632.3628 | 69.52% | 97.65% |
| **3** | 705565548 | 44937.6185 | 64.41% | 97.4% |
| **4** | 776149682 | 38768.7154 | 65.55% | 97.58% |
| **5** | 488335391 | 52050.2442 | 72.57% | 97.63% |
| **6** | 461509960 | 32169.9401 | 60.36% | 97.38% |
| **mean** | 637281864.8 | 40732.11063 | 66.91% | 97.51% |
| **stdev** | 118797857.5 | 6759.561304 | 0.039671442 | 0.001170114 |

**Supplementary Table2 Summary of single-cell ATAC sequencing (scATAC-seq) data.** Summarizes the scATAC-seq data for six samples, showing an average of 637.3 million total ATAC reads, 40,732 mean reads per cell, 66.91% sequencing saturation, and 97.51% valid barcode rates.

| **cell type** | **comparison group** | **regulation** | **DEG number** | **DAP number** | **DEG-DAP pair (direct integration)** | **DEG-DAP pair (TF integration)** |
| --- | --- | --- | --- | --- | --- | --- |
| **AT1** | flu | down | 1,374 | 338 | 81 | 37 |
|  | flu | up | 2,058 | 530 | 120 | 68 |
|  | hyperoxia | down | 1,095 | 347 | 62 | 55 |
|  | hyperoxia | up | 1,747 | 414 | 100 | 50 |
| **AT2** | flu | down | 2,253 | 565 | 164 | 59 |
|  | flu | up | 2,242 | 1,306 | 369 | 217 |
|  | hyperoxia | down | 1,145 | 36 | 11 | 3 |
|  | hyperoxia | up | 2,229 | 414 | 100 | 30 |
| **Ciliated** | flu | down | 1,975 | 2,353 | 480 | 162 |
|  | flu | up | 3,662 | 1,255 | 474 | 225 |
|  | hyperoxia | down | 1,695 | 346 | 66 | 12 |
|  | hyperoxia | up | 2,971 | 2,669 | 571 | 70 |
| **Secretory** | flu | down | 1,240 | 1,299 | 218 | 91 |
|  | flu | up | 2,237 | 485 | 111 | 63 |
|  | hyperoxia | down | 1,513 | 434 | 77 | 9 |
|  | hyperoxia | up | 2,553 | 1,776 | 428 | 76 |

**Supplementary Table 3 Summary of integrating single-cell RNA sequencing (scRNA-seq) and single-cell ATAC sequencing (scATAC-seq).** Summarize the number of differentially expressed genes (DEGs) and differentially accessible peaks (DAPs) and their integration pairs identified through both direct integration and transcription factor (TF) integration across various pulmonary cell types (AT1, AT2, Ciliated, and Secretory) under influenza (flu) and hyperoxia conditions, detailing both upregulated and downregulated regulations for each comparison group.

|  | **AT1** | **AT2** | **Ciliated** | **Secretory** |
| --- | --- | --- | --- | --- |
| **Downregulated genes in smoking** | 61 | 49 | 153 | 61 |
| **Upregulated genes in smoking** | 92 | 69 | 225 | 67 |
| **Downregulated Peaks in smoking** | 159 | 155 | 197 | 313 |
| **Upregulated Peaks in smoking** | 361 | 280 | 1651 | 723 |
| **Downregulated DEG-DAP Pair in smoking**  **(Direct integration / TF integration)** | 0 / 1 | 0 / 0 | 3 / 0 | 5 / 2 |
| **Upregulated DEG-DAP Pair in smoking**  **(Direct integration / TF integration)** | 3 / 3 | 7 / 4 | 25 / 8 | 8 / 6 |

**Supplementary Table 4 Summary of** **differentially expressed gene (DEG), differentially accessible peak (DAP), differentially expressed gene-differentially accessible peak (DEG-DAP) pair number in smoking.**

|  |  | **AT1** | **AT2** | **Ciliated** | **Secretory** |
| --- | --- | --- | --- | --- | --- |
| **DEG** | **Smoking enhances upregulated genes in flu** | 46 | 126 | 226 | 89 |
|  | **Smoking enhances downregulated genes in flu** | 235 | 489 | 297 | 137 |
|  | **Smoking reduces upregulated genes in flu** | 188 | 306 | 461 | 166 |
|  | **Smoking reduces downregulated genes in flu** | 36 | 175 | 247 | 42 |
| **DAP** | **Smoking enhances upregulated genes in flu** | 3 | 80 | 207 | 5 |
|  | **Smoking enhances downregulated genes in flu** | 136 | 379 | 469 | 283 |
|  | **Smoking reduces upregulated genes in flu** | 34 | 107 | 65 | 53 |
|  | **Smoking reduces downregulated genes in flu** | 3 | 43 | 299 | 4 |
| **DEG-DAP pair**  **(Direct integration / TF integration)** | **Smoking enhances upregulated genes in flu** | 6 / 4 | 78 / 41 | 76 / 52 | 7 / 13 |
|  | **Smoking enhances downregulated genes in flu** | 20 / 11 | 145 / 106 | 65 / 58 | 12 / 5 |
|  | **Smoking reduces upregulated genes in flu** | 2 / 10 | 87 / 281 | 64 / 229 | 5 / 0 |
|  | **Smoking reduces downregulated genes in flu** | 2 / 1 | 57 / 19 | 56 / 28 | 2 / 0 |

**Supplementary Table 5 Summary of differentially expressed gene (DEG), differentially accessible peak (DAP), differentially expressed gene-differentially accessible peak (DEG-DAP) number in smoking-influenza (flu) interactions.**

|  |  | **AT1** | **AT2** | **Ciliated** | **Secretory** |
| --- | --- | --- | --- | --- | --- |
| **DEG** | **Smoking enhances upregulated genes in hyperoxia** | 89 | 100 | 127 | 90 |
|  | **Smoking enhances downregulated genes in hyperoxia** | 52 | 76 | 57 | 91 |
|  | **Smoking reduces upregulated genes in hyperoxia** | 148 | 116 | 212 | 148 |
|  | **Smoking reduces downregulated genes in hyperoxia** | 51 | 44 | 97 | 131 |
| **DAP** | **Smoking enhances upregulated genes in hyperoxia** | 0 | 0 | 5 | 14 |
|  | **Smoking enhances downregulated genes in hyperoxia** | 0 | 4 | 10 | 10 |
|  | **Smoking reduces upregulated genes in hyperoxia** | 310 | 209 | 742 | 328 |
|  | **Smoking reduces downregulated genes in hyperoxia** | 153 | 22 | 66 | 91 |
| **DEG-DAP pair**  **(Direct integration / TF integration)** | **Smoking enhances upregulated genes in hyperoxia** | 0 / 0 | 0 / 0 | 1 / 0 | 3 / 0 |
|  | **Smoking enhances downregulated genes in hyperoxia** | 0 / 0 | 0 / 0 | 1 / 0 | 2 / 0 |
|  | **Smoking reduces upregulated genes in hyperoxia** | 3 / 0 | 3 / 0 | 17 / 2 | 9 / 2 |
|  | **Smoking reduces downregulated genes in hyperoxia** | 1 / 0 | 1 / 0 | 0 / 0 | 1 / 0 |

**Supplementary Table 6 Summary of differentially expressed gene (DEG), differentially accessible peak (DAP), differentially expressed gene-differentially accessible peak (DEG-DAP) pair number in smoking-hyperoxia interactions.**

| **Cell type** | **Smoking enhances upregulated TFs in flu** | **Smoking enhances downregulated TFs in flu** | **Smoking reduces upregulated TFs in flu** | **Smoking reduces downregulated TFs in flu** |
| --- | --- | --- | --- | --- |
| **AT2** | Arntl, DLX5, Dlx2, ELK1, ELK3, ERF, ERG, ETS1, ETV3, FEV | ALX3, ARGFX, ASCL1, Creb3l2, DMRT3, ELF2, ELF4, EN1, ETV6, EVX1 | GATA5, ISL2, MSX2, Nobox, PITX3, PRDM1, RUNX3, SOX13, Stat2, TCF3 | Arntl, Dux, GATA5, HOXB6, Hic1, NFIC, NR1I3, PITX3, PRDM1, TCF3 |
| **Ciliated** | KLF16, MEOX1, Mlxip, Msx3, NFYC, NKX2-3, POU3F4, POU5F1B, POU6F1, REL | Arntl, Atf1, BARHL1, Bhlha15, ELF2, ELF4, ELF5, ELK1, ELK3, ERF | HAND2, KLF11, KLF16, Mlxip, NFIX, NKX2-5, NKX6-3, Nobox, POU6F1, SOX4 | BSX, DRGX, EMX1, GSX2, HAND2, HOXA1, HOXA2, HOXB2, HOXB5, HOXD3 |

**Supplementary Table 7 Summary of top transcription factors (TFs) of smoking-flu interaction.** Key TFs were prioritized based on their TF ratio, calculated as the occurrence frequency of a specific TF after integration divided by its occurrence frequency before integration.

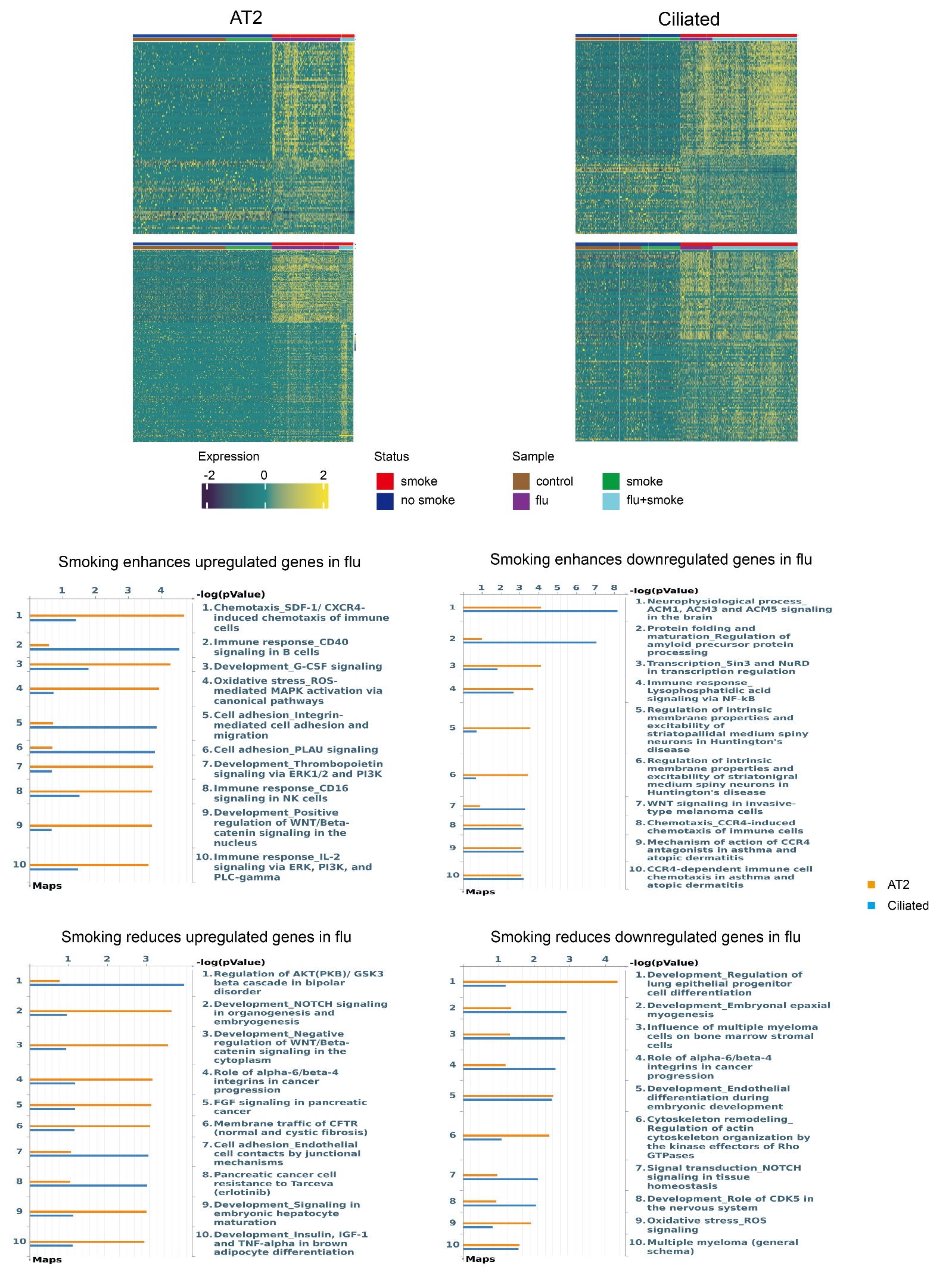

**Supplementary Fig. 1 Direct integration of single-cell RNA sequencing (scRNA-seq) and single-cell ATAC sequencing (scATAC-seq) to analyze influenza (flu) infection in response to smoking.** Top: Heatmaps showing positively interacting influenza (flu)-smoking genes in the upper panels, where smoking enhances the expression of influenza (flu)-upregulated genes or suppresses flu-downregulated genes and negatively interacting flu-smoking genes in the lower panels, where smoking suppresses influenza (flu)-upregulated genes or enhances flu-downregulated genes, across two epithelial cell types (AT2 and Ciliated). Bottom: Pathway enrichment analysis for positively and negatively smoke-influenza (flu) interacting genes.
